# Superfolder mTurquoise2^ox^ optimized for the bacterial periplasm allows high efficiency *in vivo* FRET of cell division antibiotic targets

**DOI:** 10.1101/415174

**Authors:** Nils Y. Meiresonne, Elisa Consoli, Laureen M. Y. Mertens, Anna Chertkova, Joachim Goedhart, Tanneke den Blaauwen

## Abstract

Fluorescent proteins (FP)s are of vital importance to biomedical research. Many of the currently available fluorescent proteins do not fluoresce when expressed in non-native environments, such as the bacterial periplasm. This strongly limits the options for applications that employ multiple FPs, such as multiplex imaging or FRET. To address this issue, we have engineered a new cyan fluorescent protein based on mTurquoise2 (mTq2). The new variant is dubbed superfolder turquoise 2 ox (sfTq2^ox^) and is able to withstand challenging, oxidizing environments. sfTq2^ox^ has improved folding capabilities and can be expressed in the periplasm at higher concentrations without toxicity. This was tied to the replacement of native cysteines that may otherwise form promiscuous disulfide-bonds. The improved sfTq2^ox^ has the same spectroscopic properties as mTq2, i.e. high fluorescence lifetime and quantum yield. The sfTq2^ox^-mNeongreen FRET pair allows the detection of periplasmic protein-protein interactions with energy transfer rates exceeding 40 %. Employing the new FRET pair, we show the direct interaction of two essential periplasmic cell division proteins FtsL and FtsB and disrupt it by mutations, paving the way for *in vivo* antibiotic screening.

## Introduction

In Gram-negative bacteria the cytoplasm is enveloped by an inner membrane (IM) and an asymmetric outer membrane (OM). The space between the IM and OM is called the periplasm and contains the protective peptidoglycan layer. As much as 30 % of *Escherichia coli*’s proteins are predicted to localize to the envelope and many essential processes function fully or partly within the periplasm^1^. The protein interactions in the periplasm are of great interest for biotechnological and medical purposes like synthesis of exogenous proteins and antibiotic development^2^. The most direct way to observe these proteins in living cells is by fluorescence microscopy of genetically encoded fusions to fluorescent proteins (FP)s. Fluorescence also provides a means of detecting protein-protein interactions by Förster resonance energy transfer (FRET). The stringent distance dependence for FRET is ideal to detect direct protein-protein interactions as these also occur in the nanometer range, whereas indirect protein interactions usually occur on a larger distance scale and are not detectable by FRET^3^.

*In vivo* studies of proteins in the periplasm are challenging because of its oxidizing environment and toxicity associated with protein over-expression^4^. Expression of FPs in the periplasm does not always result in fluorescence and only a limited number of FPs have been shown to fold and mature under periplasmic conditions. Recently, we observed good expression of mNeongreen (mNG) and used it as a donor to the mCherry (mCh) acceptor FP in an *in vivo* periplasmic FRET assay^4^. The mNG-mCh FRET pair has an R0 of 5.5 nm (the distance at which 50% FRET occurs) and allowed the detection of periplasmic protein-protein interactions with a dynamic range of up to 16 % energy transfer efficiency. However, this is only half the range of what can be achieved for cytoplasmic FRET pairs.

To improve our periplasmic FRET assay, popular existing FPs were screened for periplasmic fluorescence but none was adequate. Therefore, we rationally designed a novel FP dubbed sfTq2 and further optimized it for periplasmic functionality creating sfTq2^ox^. This process revealed the factors important for periplasmic fluorescence. sfTq2^ox^ has biophysical properties equal to its parent FP mTq2. Expressing it in the periplasm comes at greatly reduced toxicity, resulting in bright cyan fluorescence. sfTq2^ox^ forms a FRET-pair with mNG with an R_0_ of 6.0 nm and allows exceptionally high rates of energy transfer in the cytoplasm and periplasm of *E. coli*. Employing our new assay, we show the periplasmic interaction of the essential cell division proteins FtsB and FtsL. This work breaks ground for new research and provides microbiologists with new tools to use fluorescent techniques in the periplasm.

## Results

### Robust folding is the first prerequisite for periplasmic FP fluorescence

To optimize our periplasmic FRET assay, a higher R_0_ value FRET-pair was sought for. The periplasm is an oxidative environment and cysteines may form promiscuous disulfide bridges resulting in non-fluorescent oligomers^5^. The *Anthozoa* derived mFruits and mScarlets were thought to be good candidate acceptor FPs because of their lack of native cysteines and favorable spectroscopic properties to form a FRET-pair with the established^4^ periplasmic FP mNG instead of mCherry (**Table S1**). The FPs were expressed in the periplasm of *E. coli* by co-translational translocation through the sec-translocase (**Fig. 1**). Unfortunately, they were unable to fold or mature quickly (**Supplementary text 1 and figures**). A clear exception was mScarlet-I, which was able to fluoresce quite well in the periplasm. Still, mCherry produced stronger signals despite its modest spectroscopic properties (**Fig. S3**). These results show that the periplasmic conditions can be inhospitable for FPs regardless of their lack of cysteines (or high R_0_ FRET-pairing) and suggest that their proper folding is the first prerequisite.

**Figure 1.**
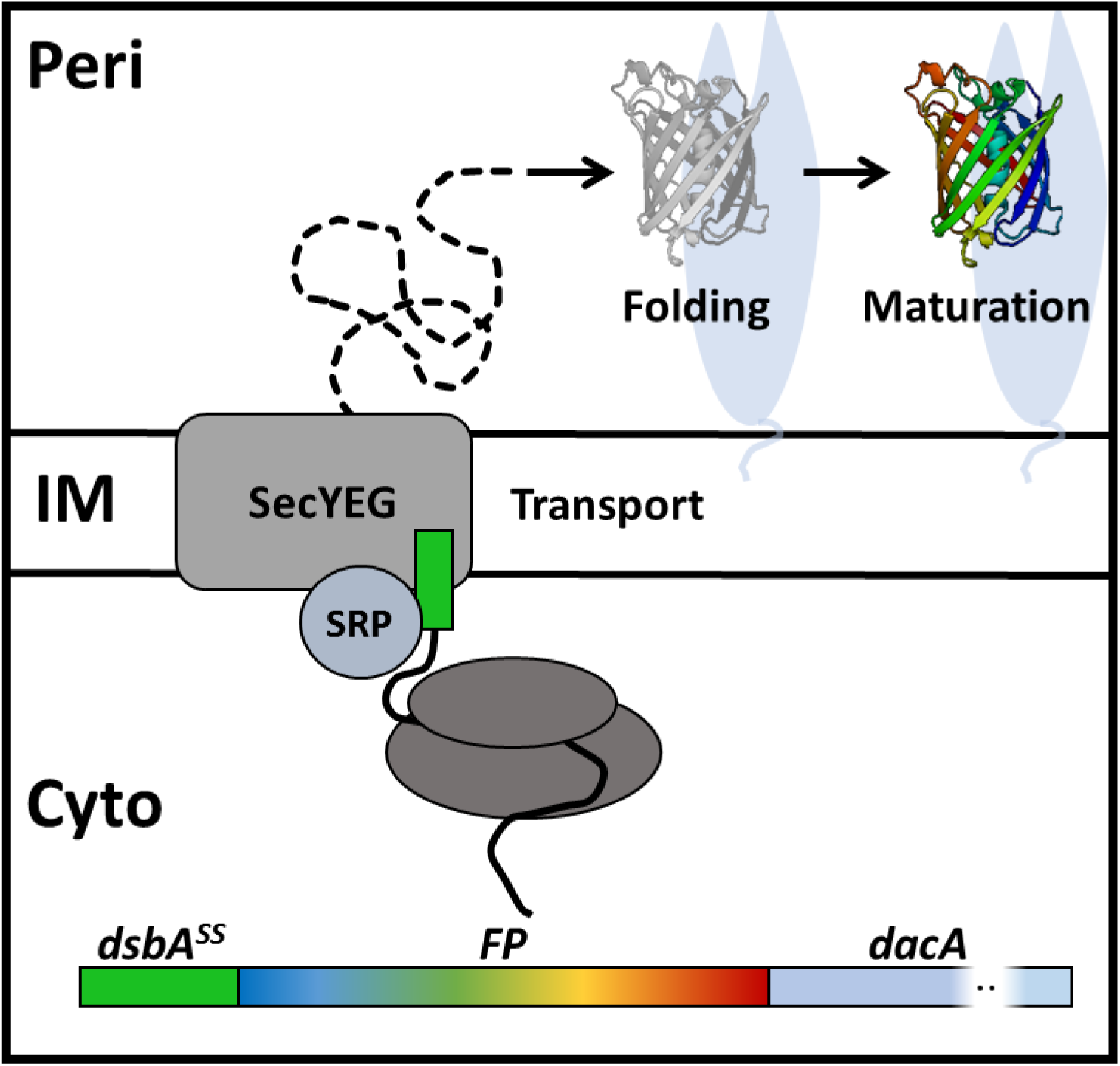
Co-translational expression of FPs in the periplasm. FP fusions were expressed in the periplasm and attached to the periplasmic (PERI) side of the inner membrane (IM) through PBP5 (encoded by *dacA)*. The DsbA signal sequence is a substrate for the signal recognition particle (SRP) that directs co-translational transport through the SecYEG translocase of the N-terminal fused gene product translated in the cytoplasm (CYTO). The FP-fusion is translocated to the periplasm and the signal sequence is cleaved off. Here the FP is exposed to periplasmic conditions under which it needs to efficiently fold its ß-barrel and subsequently mature its chromophore to become fluorescent. FPs that are unable to fluoresce in the periplasm thus have problems with either folding or chromophore maturation.

### Superfolder mTq2 folds and matures in the periplasm

*Aequorea victoria* derived FPs contain two cysteines, C48 and C70, and were therefore initially not tested. However, sfGFP folds and fluoresces in the periplasm of *E. coli*^6^ despite the presence of cysteines^7^. This observation further suggests that folding rate may be more important than the presence of cysteines for optimal periplasmic chromophore development.

Bright cyan FPs are great FRET donors because of their high quantum yield (QY). mTurquoise2 (mTq2) has a QY of 93 % and was shown to allow high FRET efficiencies with mNG^8–10^. It can be paired with green, yellow or orange acceptors that have a high molar extinction coefficient (ε) resulting in relatively high R_0_ values and strong FRET-pairs. mNG is the preferred FRET acceptor of mTq2 in eukaryotic cells with an R_0_ of 6.0 nm and FRET efficiencies ranging up to 50-60 %^9,10^.

mTq2 was tested in the periplasm of *E. coli* but it did not fluoresce (**Fig. 2a**). Since mTq2 is also derived from *Aequorea victoria,* super folding mutations S30R, Y39N, N105T F99S, M153T, V163A and I171V^7,11^ were introduced based on sfGFP to create superfolder mTq2 (sfTq2). sfGFP mutation Y145F was not included since it is close to the chromophore and could affect maturation and indeed negatively impacts periplasmic fluorescence (**Fig. S4**). A206V was not introduced since it may increase dimerization tendency^12^.

**Figure 2.**
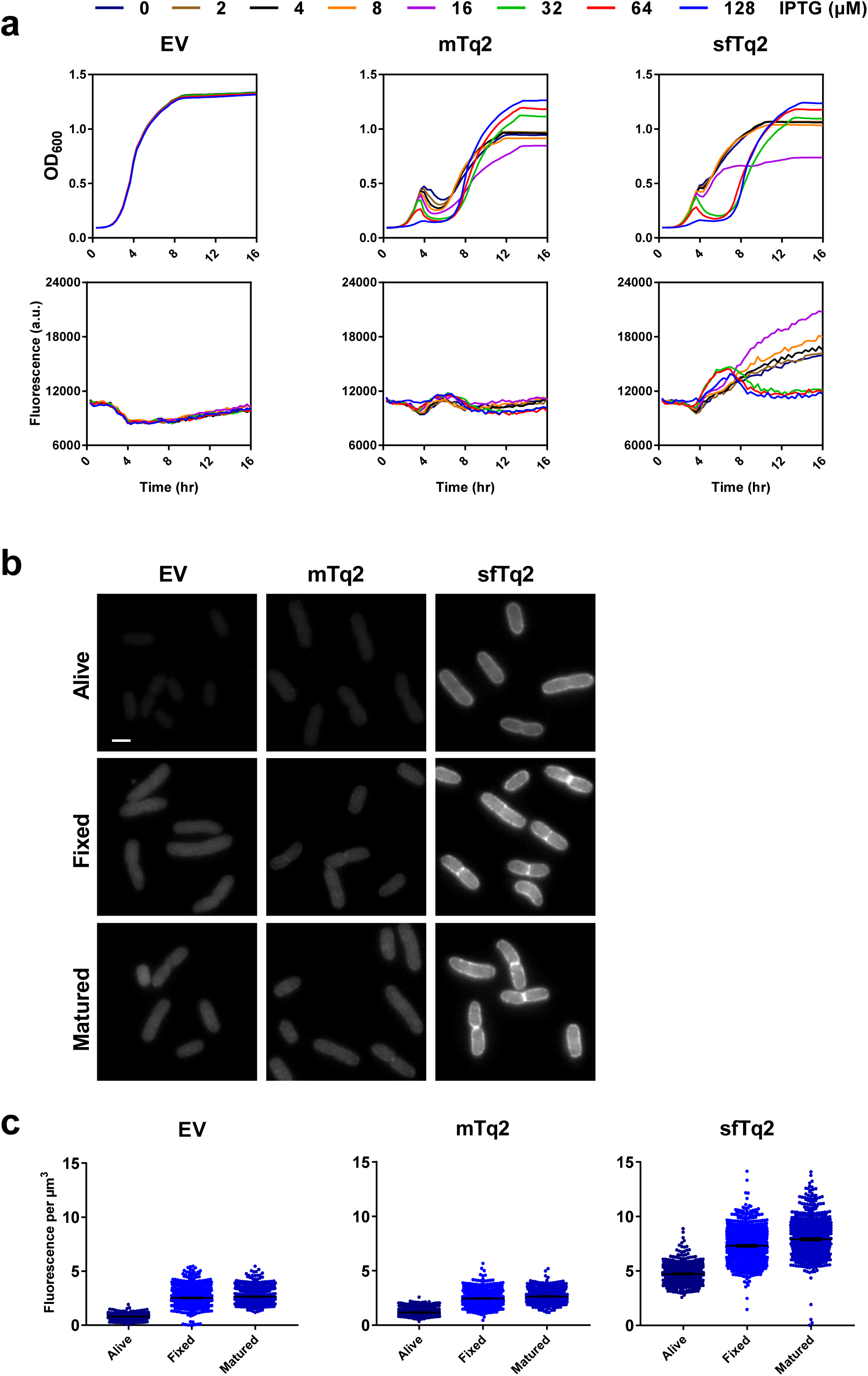
Superfolder mTurquoise2 fluoresces in the periplasm. a) Plate reader growth data in rich medium at 37 °C show that periplasmic sfTq2 expression is less toxic at high induction than the original mTq2. Plate reader fluorescence measurements confirm that periplasmic sfTq2 gives fluorescent signals while mTq2 does not. b) Fluorescence microscopy of living cells expressing an empty vector control, mTq2-PBP5 or sfTq2-PBP5 reveals fluorescence only from the sfTq2 fusion. Fixation of the same cells does not result in a decreased sfTq2 signal. Overnight maturation of the fixed cells at RT to allow for possible chromophore (re)formation (indicated as “matured”) did not result in an increase of fluorescence for periplasmic mTq2 suggesting it had not folded properly while sfTq2 showed a strong periplasmic signal. All photographs are shown with the same gray values (100-4000) for comparison and the scale bar represents 2 µm. c) Quantification of the control, mTq2 and sfTq2 cultures for the living, fixed and matured cells shows no difference between the empty vector control and the mTq2 cells. The sfTq2 cells showed a small maturation effect. The error bars at the mean represent the 95 % confidence interval. The number of cells measured were: Alive) EV = 1018, mTq2 = 650 and sfTq2 =571. Fixed) EV = 1176, mTq2 = 1046, sfTq2 = 884. Fixed and matured) EV = 1142, mTq2 = 931, sfTq2 = 988.

mTq2 and sfTq2 fusions were expressed in the periplasm of *E. coli* growing in a plate reader at a concentration range of the inducer IPTG (**material and methods**). Cultures expressing either mTq2 or sfTq2 resulted in toxicity correlated with induction concentrations previously reported with other FPs^4^. However, sfTq2 could be expressed at higher levels with less toxicity compared to mTq2 supporting the notion that fast folding improves expression. Strikingly, the sfTq2 fusion resulted in periplasmic fluorescence whereas the mTq2 fusions did not (**Fig. 2a**). This shows that the superfolder mutations enable sfTq2, like sfGFP, to fold and mature in the periplasm.

Cells expressing mTq2 and sfTq2 in the periplasm at non-toxic concentrations were imaged by fluorescence microscopy. Quantification of the signals from living, fixed and fixed and matured samples showed periplasmic fluorescence of sfTq2 only (**Fig. 2bc**). Western blot analysis confirmed the similar expression levels of mTq2 and sfTq2 (**shown in the next section**).

### Development and optimization of sfTq2^ox^

Superfolder mutations allowed sfTq2 to fold and mature in the periplasm. Next, we tested how the new donor performed in FRET experiments with mNG. Cytoplasmic mNG-sfTq2 is capable of the same rates of energy transfer as mNG-mTq2 with *Ef*_*A*_ values of 64.5 ± 3.0 % and 61.3 ± 6.4 %, respectively. However, periplasmic mNG-sfTq2 only amounted to 18.8 ± 2.1 % (**Table 1**). Possibly, the superfolder mutations in sfTq2 did not fully protect its cysteines from forming non-fluorescent hetero- and oligomers in the periplasm (**Supplementary text 2 and figures**).

**Table 1.** FRET efficiencies obtained with mTq2 variants measured by fluorometry.

| Cytoplasmic FRET | Donor FP to mNG | Fusion | $Ef_A$ (%) | SD | Repeats |
| --- | --- | --- | --- | --- | --- |
| **Positive control tandems** |  |  |  |  |  |
| mNG-mTq2 | mTq2 | Free floating | 61.3 | 6.4 | 4 |
| mNG-sfTq2 | sfTq2 | Free floating | 64.5 | 3.0 | 7 |
| mNG-sfTq2<sup>C70V</sup> | sfTq2<sup>ox</sup> | Free floating | 66.4 | 3.1 | 8 |
| Periplasmic FRET |  |  |  |  |  |
| **Positive control tandems** |  |  |  |  |  |
| mNG-sfTq2 | sfTq2 | OmpA177 | 18.8 | 2.1 | 6 |
| mNG-sfTq2<sup>C48S</sup> | C48S | OmpA177 | 32.9 | 1.9 | 2 |
| mNG-sfTq2<sup>C70V</sup> | sfTq2<sup>ox</sup> | OmpA177 | 39.1 | 3.8 | 11 |
| mNG-sfTq2<sup>C48S-C70V</sup> | C48S-C70V | OmpA177 | 40.3 | 1.2 | 2 |
| mNG-sfTq2<sup>C70V</sup> | sfTq2<sup>ox</sup> | OmpA177* | 42.5 | 3.1 | 6 |
| mNG-sfTq2<sup>C70V</sup> | sfTq2<sup>ox</sup> | LpoB73 | 39.8 | 2.1 | 5 |
| mNG-sfTq2<sup>C70V</sup> | sfTq2<sup>ox</sup> | NlpA<sup>ss</sup> | 41.2 | 2.4 | 7 |
| mNG-sfTq2<sup>C70V</sup> | sfTq2<sup>ox</sup> | MalF<sup>ss-mss</sup> | 42.9 | 2.5 | 3 |
| **Negative controls** |  |  |  |  |  |
| mNG + sfTq2 | sfTq2 | IM-OM | -1.4 | 1.4 | 4 |
| mNG + sfTq2<sup>C70V</sup> | sfTq2<sup>ox</sup> | IM-OM | -1.3 | 2.2 | 13 |
| mNG + sfTq2<sup>C70V</sup> | sfTq2<sup>ox</sup> | IM-IM | -0.7 | 1.7 | 10 |
| **Biological interactions** |  |  |  |  |  |
| PBP5 + PBP5 | sfTq2 | IM-IM | 4.2 | 2.7 | 5 |
| PBP5<sup>S44G</sup> + PBP5<sup>S44G</sup> | sfTq2 | IM-IM | 8.8 | 4.2 | 6 |
| FtsB + FtsL | sfTq2<sup>ox</sup> | IM-IM | 19.2 | 4.5 | 8 |
| FtsL + FtsB | sfTq2<sup>ox</sup> | IM-IM | 15.6 | 2.6 | 8 |
| FtsL<sup>m4</sup> + FtsB | sfTq2<sup>ox</sup> | IM-IM | 3.1 | 2.1 | 3 |
| FtsL<sup>m4</sup> + FtsB<sup>m4</sup> | sfTq2<sup>ox</sup> | IM-IM | 3.0 | 3.4 | 3 |

<sup>SS</sup> is signal sequence, <sup>mss</sup> membrane spanning sequence, LpoB73 indicates residue 1-73 of LpoB, OmpA177 indicates residue 1-177 of OmpA, \* indicates a different linker between mNG and sfTq2<sup>ox</sup> (EF instead of EL due to differences in cloning). **Table S2** shows the FRET efficiencies for the mTq2 variants measured with the plate reader. Representative unmixing data is shown in **Fig. S16**.

To further improve the expression of sfTq2 in the periplasm, site-directed mutagenesis was performed to replace sfTq2’s C48 with serine and C70 with serine, methionine or valine. Serine closely resembles cysteine with a hydroxyl-instead of a sulfhydryl group. Methionine also contains a sulfur that is not nucleophilic and therefore should not participate in disulfide bond formation. Valine should resemble the more hydrophobic state of cysteine better than serine and was shown to aid fluorescence of oxBFP in the eukaryotic ER^5^.

Single and double cysteine mutants of sfTq2 performed better than sfTq2 in the periplasm with reduced toxicity and improved fluorescence (**Fig. 3, Fig. S6**). They were expressed as protein fusions in the periplasm of *E. coli* under non-toxic conditions and the fluorescence signals of living, fixed and fixed and matured samples were quantified. sfTq2 mutants C48S, C70S, C70V and C48S-C70V showed much brighter signals from the periplasm (**Fig. 3bc**). The methionine sfTq2 variants did not perform better than sfTq2 in terms of fluorescence (**Fig. S6**). Western blotting showed that all cysteine replacement mutants were better expressed in the periplasm regardless of fluorescence signals (**Fig. 3d**). This is in line with the secretory improvement of GFP to the eukaryotic ER by cysteine replacements^13^. The enhanced periplasmic production but dim fluorescence of some sfTq2 variants suggests poor chromophore maturation.

**Figure 3.**
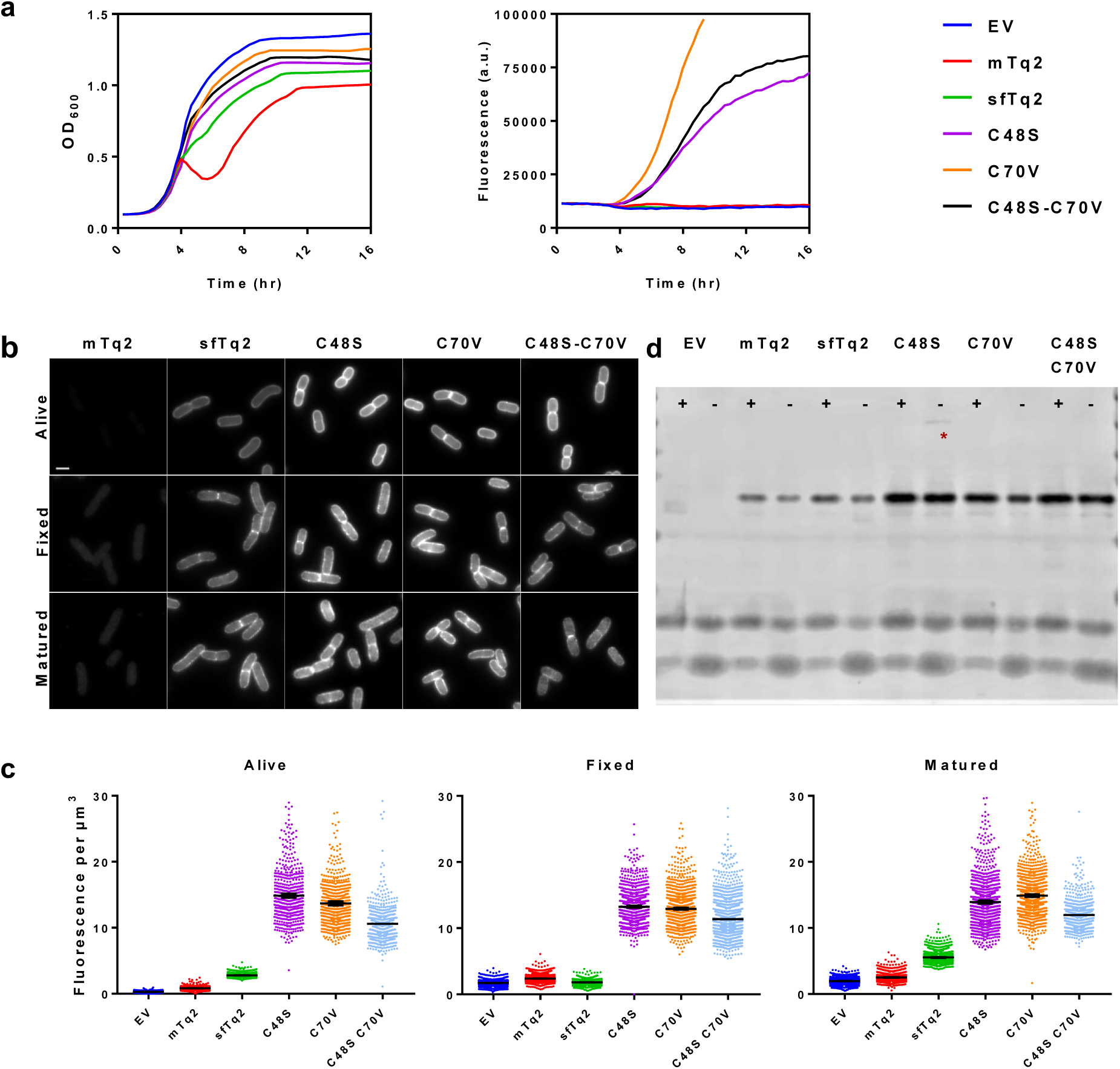
Cysteine-replaced sfTq2 variants perform better in the periplasm. a) Periplasmic expression of sfTq2-PBP5 variants in LMC500 grown in rich medium at 37°C at relatively non-toxic induction conditions results in large differences in cyan fluorescence. The C70V variant performs better than C48S and the double cysteine mutant version. b) Microscopy of LMC500 grown in rich medium and induced with 15 µM IPTG shows strong fluorescence for the sfTq2-PBP5 variants in the periplasm. For comparison the grayscale of all photographs are the same (80-7000) and the scale bar represents 2 µm. c) Quantification of the images confirms bright fluorescence signals from the single C48S and C70V sfTq2 variants. Differences in fluorescence with the prolonged plate reader induction suggest that folding difficulties may still be a problem. The error bars at the mean represent the 95 % confidence interval. The number of cells measured were between: Alive 500-1000, Fixed 1000-1500 and Matured 1000-2000 except for EV-Alive n = 327 and C48S-C70V-Fixed n = 775. d) mTq2 is produced at similar levels as sfTq2 but does not fold properly, cysteine mutants of sfTq2 are produced at higher levels. Anti-GFP immunoblotting of the corresponding samples shows that the fusion proteins are intact and that C48S has a minor propensity to form higher order complexes (asterisk). The + and – signs indicate the presence or absence of reducing agent DTT in the sample, respectively.

Site directed random mutagenesis of sfTq2 cysteines revealed that residue 48 does not allow much variation besides the original cysteine or serine while a variety of amino acids are accepted as residue 70. Several brightly fluorescing sfTq2 variants were found but none of them showed improved periplasmic fluorescence as compared to C70V (**Supplementary text 3, Fig. S7**). Cysteine mutants in the parental mTq2 did not increase periplasmic fluorescence although a reduction of expression toxicity was observed (**Fig. S8**). sfGFP mutant sfGFP^C70V^ showed similar periplasmic fluorescence compared to its predecessor and expression was slightly less toxic (**Fig. S9**). This shows that cysteines are involved in translocation toxicity and subsequent low production of periplasmic fusion proteins.

All experiments showed that sfTq2^C70V^ was the best periplasmic variant to be used in the bacterial periplasm in terms of reduced toxicity, fluorescence intensity and fast folding or maturation. sfTq2^C70V^ was named sfTq2^ox^. Subsequent comparisons of sfTq2 and sfTq2^ox^ as periplasmic fusions to several IM and OM localized proteins confirmed the superior performance of sfTq2^ox^ (**Figs. S10-12**).

### The superior periplasmic properties of sfTq2^ox^ come without a trade-off

Introducing superfolder mutations in mTq2 allowed fluorescence in the periplasm of *E. coli*. The additional C70V mutation in sfTq2^ox^ resulted in a reduction of expression toxicity and the brightest periplasmic fluorescence. Interestingly, mTq2, sfTq2 and sfTq2^ox^ were equally bright when expressed in the cytoplasm (**Fig. 4d**, **Fig. S13**). To verify the *in vivo* spectroscopic properties of the mTq2 variants the fluorescence lifetime and cellular brightness were determined.

**Figure 4.**
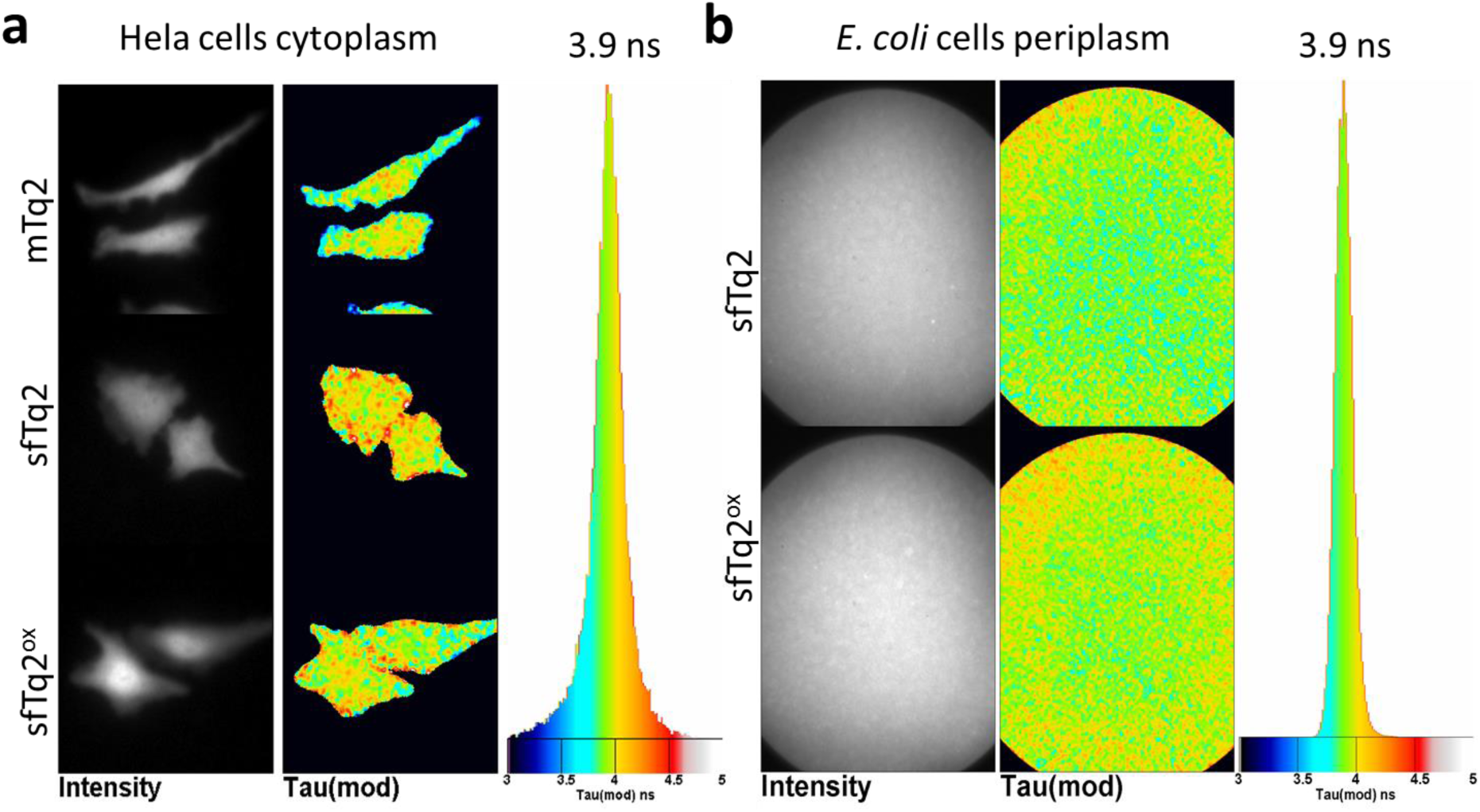
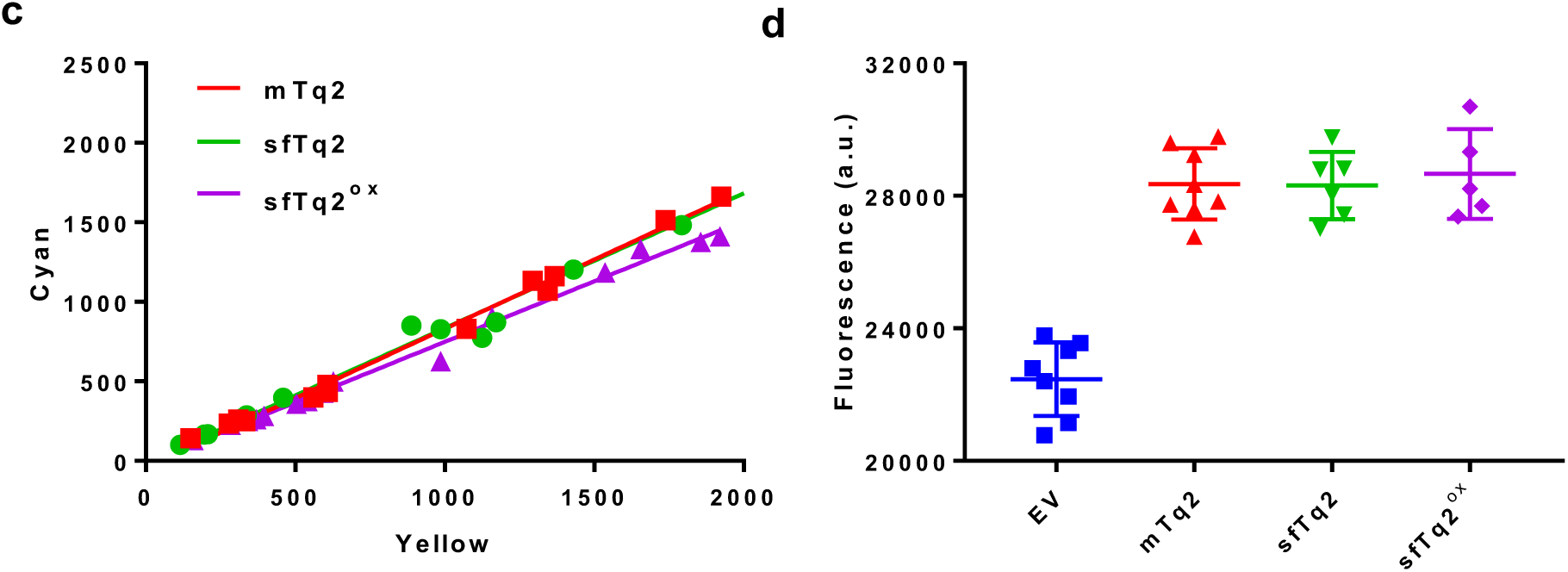
sfTq2^ox^ comes without a trade-off. a) Frequency domain FLIM of HeLa cells expressing mTq2, sfTq2 or sfTq2^ox^ in the cytoplasm reveals a similar average fluorescence lifetime of 3.9 ns. b) FLIM of a suspension of *E. coli* cells expressing periplasmic OmpA-sfTq2 or OmpA-sfTq2^ox^ also show a similar fluorescence lifetime of 3.9 ns. c) The cellular brightness of mTq2, sfTq2 and sfTq2^ox^ is equal in Hela cells. The slope coefficients of the fits are, respectively: 0.87, 0.84 and 0.76. d) Fixed samples of LMC500 cells at OD_450_ = 1.00 expressing cytoplasmic mTq2, sfTq2 or sfTq2^ox^ show equal amounts of fluorescence.

The quantum yield of cyan FPs is directly related to their fluorescence lifetime^8,14^. Therefore, the lifetimes of mTq2 and sfTq2 and sfTq2^ox^ were measured from the cytoplasm of eukaryotic (HeLa) cells using frequency domain fluorescence lifetime imaging microscopy (FLIM). This resulted in an identical distribution of lifetimes with an average of 3.9 ± 0.1 ns suggesting no differences in QY between the three variants (**Fig. 4a**). The fluorescence lifetime of sfTq2 and sfTq2^ox^ expressed in the periplasm also showed a similar average lifetime of 3.9 ± 0.1 ns showing that periplasmic conditions did not alter their lifetime and suggesting an unaltered QY (**Fig. 4b**). Agar colonies of *E. coli* expressing cytoplasmic mTq2, sfTq2 or sfTq2^ox^ were imaged and gave a similar average lifetime of 3.8 ± 0.2 ns (**Fig. S13**).

To analyze the cellular brightness of sfTq2 and sfTq2^ox^ self-cleaving viral peptide (2A) linked tandems with super yellow FP 2 (sYFP2) were expressed in HeLa cells^10^. The equal expression of both FPs^15^ showed similar correlation for all three mTq2 variants reflecting an equal cellular brightness (**Fig. 4c**). Quantification of mTq2, sfTq2 and sfTq2^ox^ expressed in the cytoplasm of *E. coli* further supports this (**Fig. 4d, Fig. S13**). Having determined that the lifetime and cellular brightness of sfTq2^ox^ are the same as that of mTq2 and sfTq2, it is reasonable to assume that the extinction coefficient is also similar. The unaltered lifetime also implies that the spectroscopic properties (quantum yield) of mTq2 may be used for calculating R_0_ and the quantification of *Ef*_*A*_ values.

mTq2 is monomeric^12^ and will therefore not obscure FRET experiments by dimerization or oligomerization. The superfolder and C70V mutations may have impacted this property of mTq2. Therefore, the tendency of sfTq2 and sfTq2^ox^ to form undesired oligomers was assessed by OSER assay^16^ showing equal results for both proteins, suggesting the same low propensity to aggregate as mTq2 (**Fig. S14**).

Further confirmation that sfTq2 and sfTq2^ox^ have the same spectral properties as mTq2 comes from their FRET efficiencies in the cytoplasm. Tandem fusions of mNG with mTq2, sfTq2 or sfTq2^ox^ resulted in significantly similar *Ef*_*A*_ values of 61 ± 6 %, 65 ± 3 % and 66 ± 3 %, respectively (**Table 1**). Interestingly, the same constructs expressed in agar colonies of *E. coli* gave an average lifetime of 2.5 ± 0.1 ns, a 35 % reduction in fluorescence lifetime compared to their single donor FPs (**Fig. S13**). This is similar to the 33 % energy transfer observed for mNG-mTq2 in eukaryotic cells that were measured using the same technique^10^.

Taken together, these results strongly indicate that the engineering of sfTq2^ox^ comes without a trade-off compared to its parent mTq2 with the added benefits for periplasmic expression, bright fluorescence and reduced toxicity.

### High dynamic range periplasmic FRET with sfTq2^ox^

mTq2 forms a good FRET-pair with mNG with an R_0_ of 6.0 nm. The cytoplasmic tandems of mTq2 and sfTq2 gave similar rates of energy transfer. Yet, the periplasmic mNG-sfTq2 tandem gave an *Ef*_*A*_ value of 18.8 ± 2.1 % suggesting differences in terms of periplasmic functionality. The bright periplasmic fluorescence of cysteine-replaced sfTq2 variants suggested that they would make better donors to mNG for periplasmic FRET. mNG-sfTq2 tandems containing the cysteine mutants, associated with the OM through OmpA, were tested for their *in vivo* FRET efficiencies. This indeed greatly improved *Ef*_*A*_ values to 32.3 ± 1.9, 42.5 ± 3.1 % and 40.3 ± 1.2 % for sfTq2^C48S^, sfTq2^ox^ or sfTq2^C48S-C70V^, respectively. No degradation or cleavage products were detected by western blotting (**Fig. S15**).

Since sfTq2^ox^ showed the best periplasmic behavior compared to all other cysteine mutant variants, mNG-sfTq2^ox^ fusions were made to other periplasmic proteins to exclude localization effects. Tandems were associated with the OM or IM through the translocation and lipidation signals of lipoproteins LpoB and NlpA, respectively. A third tandem was localized to the IM through the first membrane spanning sequence of MalF (**Fig. 5**). All periplasmic tandems resulted in *Ef*_*A*_ values of 40 % (**Table 1**) suggesting that this is the highest attainable FRET efficiency for mNG-sfTq2^ox^ in the periplasm regardless the location of the fusion. Negative controls, assaying energy transfer by crowding conditions in the IM or between the IM and OM resulted in *Ef*_*A*_ values of −0.7 ± 1.7 % and −1.3 ± 2.2 %, respectively (**Fig. 5**). All assayed *Ef*_*A*_ values are shown in **Table 1**. The measured FRET efficiencies were confirmed using a separate 96 wells plate reader set up showing that the throughput of FRET assays using the sfTq2^ox^-mNG pair can be greatly increased (**Table S2**).

**Figure 5.**
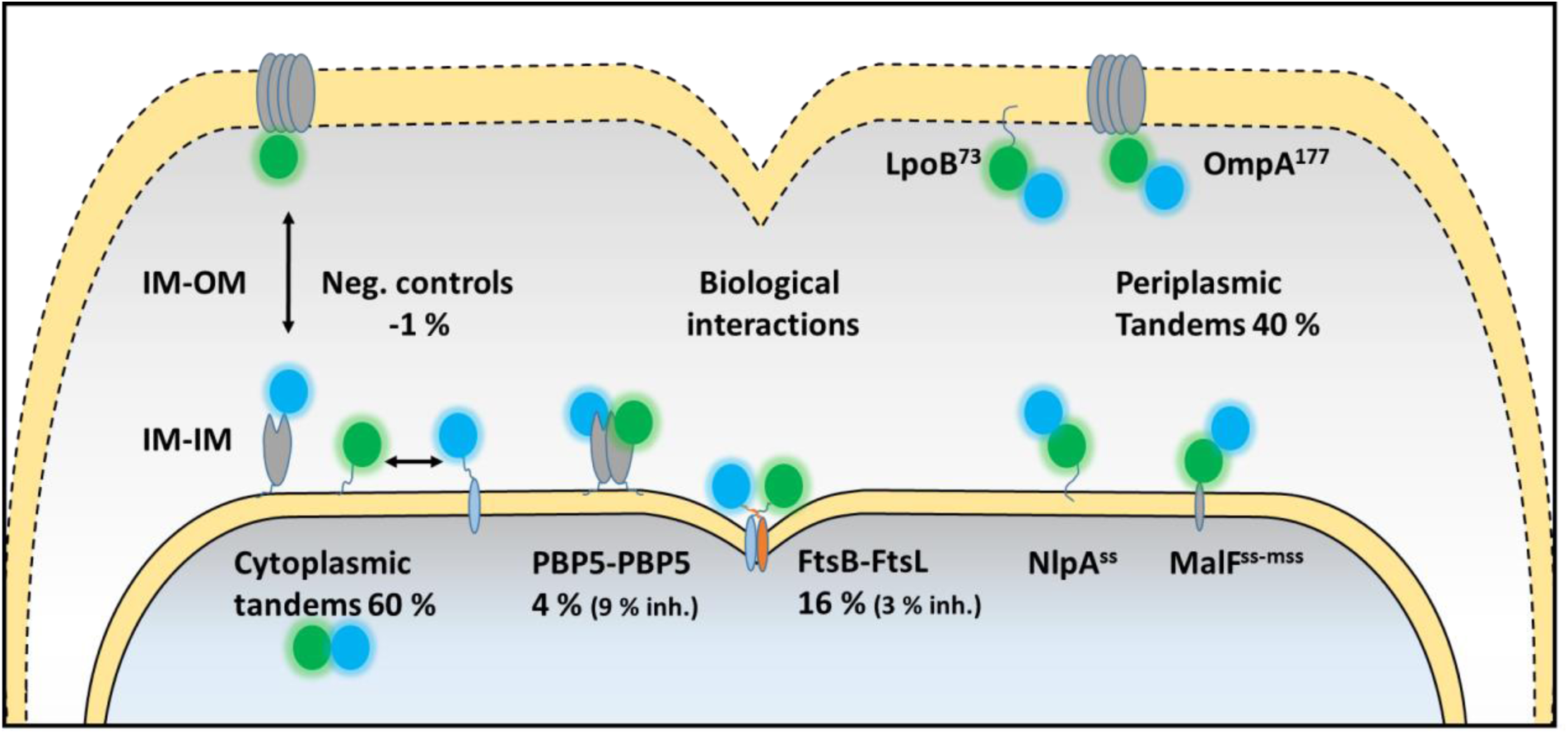
Schematic overview showing the subcellular localization of the periplasmic protein interactions and controls assayed with the sfTq2^ox^-mNG FRET pair. PBP5 dimerization interactions were assayed with sfTq2-mNG and are described in **supplementary text 2**.

### *In vivo* interactions of putative antibiotic targets in the periplasm

With the periplasmic sfTq2^ox^-mNG FRET-pair established, biological interactions could be assessed. The periplasm contains many relevant potential antibiotic targets including the cell division subcomplex of FtsQ, FtsL, and FtsB (FtsQLB)^17^. All three components are essential and are thought to form the regulating link between cytoplasmic and periplasmic machineries of cell division^18^. Assaying the interactions in this complex could allow for the screening of novel antibiotics *in vivo*, determining efficacy and specificity at the same time. FRET experiments assaying the interaction between FtsB-sfTq2^ox^ and FtsL-mNG or FtsB-mNG and FtsL-sfTq2^ox^ resulted in *Ef*_*A*_ values of 19.2 ± 4.5 % and 15.6 ± 2.6 %, respectively (**Table 1, Fig. 5**). A negative control for IM-associated proteins of FtsB-sfTq2^ox^ with NlpA^ss^-mNG resulted in an *Ef*_*A*_ value of −0.7 ± 1.7 %. FtsB and FtsL were reported to interact through a leucine zipper-like motif that can be disrupted by leucine to alanine substitutions^19^. The leucine zipper mutants of FtsB, FtsB^m4^, and FtsL, FtsL^m4^, are good controls as they are able to localize at the division site and complement their respective depletion strains^19^. The interactions of wild-type FtsB-mNG or mutant FtsB^m4^-mNG with mutant FtsL^m4^-sfTq2^ox^ showed decreased *Ef*_*A*_ values of 3.1 ± 2.1 % and 3.0 ± 3.4 %, respectively. Expression without degradation of wild-type and mutant FtsB and FtsL constructs was confirmed by western blotting (**Fig. S15**). The weakening of the leucine zipper motif clearly reduces the interaction between the two proteins given the decrease in *Ef*_*A*_ values. This observed reduced affinity between the two proteins may reflect a reduction in their average interaction time. In conclusion, the new FRET pair sfTq2^ox^-mNG can be used to detect periplasmic protein interactions *in vivo* and paves the way for functional screening of essential antibiotic targets.

## Discussion

*In vivo* visualization of periplasmic proteins by fusions to FPs is difficult since most do not fold or mature due to the oxidizing environment and the necessity of transport. Additionally, over-expressing periplasmic FP fusions can lead to toxicity and counter selection^4^. The limited choice of FPs is especially problematic for applications that use multiple FPs, such as multiplex imaging and FRET studies. For FRET studies, a pair of FPs with a high R_0_ is desired. The theoretical R_0_ value, however, does not take into account the proper folding and maturation of FPs *in vivo*. Therefore, the optimal FP for experiments under specific conditions should be experimentally determined. Previously we developed a periplasmic FRET assay using mNG-mCh (R_0_ = 5.5 nm) and used it to show conformational changes of penicillin-binding proteins inactivated by mutations or antibiotic treatment^4^. mCherry has a modest quantum yield (QY) and subsequent brightness, which is suboptimal for FRET experiments that measure sensitized emission. Hence, an acceptor with a higher QY would improve the dynamic range. Therefore, the closely related mFruits were tested for periplasmic functionality without any observed improvement. The also related, newer mScarlet-I showed periplasmic fluorescence but mCherry still performed better, likely due to favorable folding and maturation capabilities (**Supplementary text 1**).

mNG has a high QY and extinction coefficient and can therefore function as a donor or acceptor FP^9^. It forms an efficient FRET-pair with mTq2 (R_0_ = 6.0 nm) and by virtue of its efficient maturation allows for high energy transfer efficiencies. For this pair, FRET is accompanied by a substantial amount of sensitized emission due to the high QY of mNG^9,10^. Since mTq2 did not fluoresce in the periplasm, we focused our efforts on the development of a donor that shows good maturation in the periplasm. sfTq2 was engineered by introducing superfolder mutations S30R, Y39N, N105T F99S, M153T, V163A and I171V based on the related sfGFP, which is fluorescent in the periplasm^6^. sfTq2 was functional in the periplasm and allowed only moderate periplasmic FRET with mNG, compared to the cytoplasm where it performed comparable to mTq2 (**Fig. 1, table 1, supplementary text 2**).

The oxidative environment of the periplasm facilitates disulfide bond formation between exposed cysteines in close proximity^6^. *Aequorea victoria* derived FPs are natively adapted for cytoplasmic conditions and their cysteines are thus prone to promiscuous disulfide bonds resulting in non-fluorescent oligomers in the periplasm. Site-directed random mutagenesis of C48 and C70 in sfTq2 resulted in greatly increased periplasmic fluorescence and reduced expression toxicity. Random mutagenesis revealed that only serines or cysteines are tolerated at position 48 while position 70 allowed more variation in amino acid replacements resulting in variable levels of periplasmic fluorescence (**Supplementary text 3**). The single C70V mutation variant, called sfTq2^ox^, was able to produce bright, fast maturing, periplasmic fluorescence and reduced translocation toxicity (**Fig. 3**). It came without an observed trade-off as the same QY and brightness values were found for mTq2 and sfTq2 (**Fig. 4**). Taken together, the development of sfTq2^ox^ shows that periplasmic fluorescence of FPs first depends on their efficient folding and then on the absence of cysteines to prevent the formation of non-fluorescent oligomers and reduce expression toxicity.

Employing the mNG-sfTq2^ox^ FRET pair in the periplasm resulted in energy transfer rates of 40 % for all localized tandems, irrespective of whether they were bound close to the IM or with some distance from the OM as with the LpoB fused tandem (**Fig. 5**). FRET efficiencies of 60 % were observed for cytoplasmic mTq2, sfTq2 or sfTq2^ox^ fusions to mNG (**Table 1**). Fluorescence lifetime measurements of the cytoplasmic tandems in bacterial colonies showed energy transfer rates of 35 % for mTq2, sfTq2 and sfTq2^ox^ (**Fig. S13**), similar to results for mNG-mTq2 in eukaryotic cells^10^. Based on the R_0_ value (6.0 nm), the FRET efficiency of 60 % and 40 % would correspond to a distance between the fluorophores of 5.6 and 6.4 nm, respectively. Both values are plausible given the minimal distance of ± 3 nm between the two chromophores of the FRET pair. Yet, it cannot be excluded that the transport to the periplasm still hampers subsequent folding of a fraction of the FPs resulting in a lower FRET efficiency.

A factor influencing FRET efficiency (and R_0_ values) is the dipole-orientation (*κ*^2^) of the donor and acceptor chromophore. This would only contribute to the FRET efficiency difference if the environment of the cytoplasm or periplasm would restrict the rotational freedom of the proteins in the tandem differently. Presently, no evidence is available that would support such a difference. The comparison of mTq2 with several acceptor FPs shows that mNG is an exceptional FRET acceptor, even compared to FPs with similar spectroscopic properties^10^. Indeed, mNG is an outlier with its chordate origin compared to the usual cnidarian-derived FPs with different sequence homologies and possibly different behaviors ^9,21^.

The ability of FPs to fold and mature under non-native conditions is a property that is of increasing interest. Co-translational expression of fusion proteins in the periplasm is a good assay to select for optimized FPs. With the development of sfTq2^ox^ the palette of periplasmic functional FPs is broadened and now includes members emitting at cyan wavelengths. This allows for the *in vivo* observation of (at least) three colors of periplasmic fusion proteins. Moreover, sfTq2^ox^ forms a FRET pair with mNG that can detect periplasmic protein interactions *in vivo* at double the detection range for periplasmic FRET assays in bacteria. This new combination is therefore the preferred FRET-pair for periplasmic as well as cytoplasmic protein-interaction assays.

Using the new periplasmic FRET-pair we have demonstrated the direct interaction and disruption of essential cell division and putative antibiotic target proteins FtsB and FtsL with high rates of energy transfer. As these experiments were done in wild-type *E. coli* the actual *Ef*_*A*_ between fluorescent protein fusions may be higher since native FtsL and FtsB compete with the fusions. Interactions of the FtsQLB complex are considered to be the link between the cytosolic and periplasmic parts of cell division setting off constriction^18^. Exactly how and in what stoichiometry the complex forms and functions is still a matter of debate^19,22–25^. Interactions between FtsB and FtsL seem to rely on their coiled-coil transmembrane helices^19,24^ while the interaction of FtsQ and FtsB requires their periplasmic soluble domains and forms independently of FtsL^23,26,27^. Our findings seem to confirm the former interaction *in vivo* and provide a tool for the further dissection of FtsQLB interactions. Protein interactions often chance significantly when the activity of one of the partners is inhibited by antibiotics ^28^. The Gram-negative periplasm facilitates a myriad of essential protein interactions waiting to be investigated *in vivo*. The development of sfTq2^ox^ paves the way for multiplex periplasmic imaging and FRET applications, enabling the screening of protein interactions and novel antibiotics that may inhibit them.

## Acknowledgements

We thank Eva Asmus and Alejandro Mónton Silva (University of Amsterdam, the Netherlands) for help with the experiments. Plasmids pCR119 and pCR124 were kind gifts from Daniel Ladant (Institut Pasteur, France). mScarlet-C1, mScarletI-C1 and mScarletH-C1 were kind gifts from Theodorus W.J. Gadella Jr. (University of Amsterdam, the Netherlands). N.Y.M. was supported by the NWO, ALW open program (822.02.019) and HFSP Program RGP0034/2013. E.C. and L.M.Y.M. were supported by the European Commission via the International Training Network Train2Target (721484).

## Contributions

N.Y.M. and T.d.B. designed the study. N.Y.M., E.C., L.M.Y.M., J.G. and A.C. conducted experiments and analyzed data. N.Y.M. and T.d.B. wrote the paper. All authors contributed to the final version of the manuscript.

The authors declare no competing financial interests.

## Materials and Methods

### Bacterial strains and culturing conditions

The *Escherichia coli* K12 strains used are presented in in (Table 2). The cells were cultured in rich medium (TY: 10 g Tryptone (Bacto laboratories, Australia), 5 g yeast extract (Duchefa, Amsterdam, The Netherlands) and 5 g NaCl (Merck, Kenilworth, NJ) per liter) supplemented with 0.5% glucose (Merck) or in glucose minimal medium (Gb1: 6.33 g K_2_HPO_4_ (Merck), 2.95 g KH_2_PO_4_ (Riedel de Haen, Seelze, Germany), 1.05 g (NH_4_)_2_SO_4_ (Sigma, St. Louis, MO), 0.10 g MgSO_4_·7H_2_O (Roth, Karlsruhe, Germany), 0.28 mg FeSO_4_·7H_2_O (Sigma), 7.1 mg Ca(NO_3_)_2_·4H_2_O (Sigma), 4 mg thiamine (Sigma), and 4 g glucose per liter, pH 7.0) at 28 °C while shaking at 205 rpm. For growth in Gb1 of LMC500 and CS109 based strains 50 mg/l lysine (Sigma) was added. Growth in rich and poor medium was at 37 and 28 °C, respectively. Expression of constructs was induced with 15 µM isopropyl β-D-1-thiogalactopyranoside (IPTG, Promega, Madison WI) unless stated otherwise. Plasmids were maintained in the strains by addition of 100 µg/ml ampicillin (Sigma) or 25 µg/ml chloramphenicol (Sigma). Growth was measured by absorbance at 600 or 450 nm with a Biochrom Libra S70 spectrophotometer (Harvard Biosciences, Holliston, MA) for TY or Gb1 cultures, respectively. Fixation was done with a final concentration of 2.8% formaldehyde and 0.04% glutaraldehyde in the shaking water bath for 15 min, after which the cells were harvested. After fixation the cells were washed three times with 1 ml PBS.

**Table 2.** Strains used in this study.

| Strain | Relevant characteristics | Ref. |
| --- | --- | --- |
| DH5α | F-, supE44, hsdR17, recA1, endA1, gyrA96, thi1, relA1 | 29 |
| LMC500 | MC4100, F-, araD139, Δ(argF-lac)U169, deoC1, flbB5301, lysA1, ptsF25, rbsR, relA1, rpsL150 | 30 |
| CS12-7 | CS109 ΔdacA | 31 |

### Plasmid construction

An overview of the plasmids used in this study and their cloning strategies is given in **table S3**. The sfTq2 nucleotide sequence (encoding mTurquoise2 with S30R, Y39N, F99S, N105T and I171V) was obtained by gene synthesis (MWG biotech). New constructs were created by restriction-ligation cloning or site-directed mutagenesis. Inserted fragments were amplified by PCR from plasmid or the *E. coli* (LMC500) chromosome template, purified and restricted. For mutagenesis, a whole plasmid template was amplified by PCR using primers that contain the desired mutation. The resulting product was treated with *DpnI* to digest the methylated template plasmid. All restriction enzymes used were purchased from New England Biolabs Inc. (Ipswich, MA). For all PCR amplifications for cloning, the high-fidelity polymerase *pfu*x7 was used^32^ as per the manufacturer’s instructions. An overview of the used primers is shown in **table S4**.

### Imaging and image analysis

For imaging the cells were immobilized on 1 % agarose in water slabs on object glasses as described^33^ and photographed with a Hamamatsu ORCA-Flash-4.0LT (Hamamatsu, Naka-ku, Japan) CMOS camera mounted on an Olympus BX-60 fluorescence microscope (Tokyo, Japan) through a UPlanApo 100x/N.A. 1.35 oil Iris Ph3 objective. Images were acquired using the Micro Manager 1.4 plugin for ImageJ^34^. In all experiments, the cells were first photographed in phase contrast mode and then in fluorescence mode. The fluorescence filter cubes used were: U-MNG (red and orange, ex560/40, dic585LP, em630/75), EN-GFP (green, ex470/40, dic495LP, em525/50) and Cyan-GFPv2 (cyan, ex436/20, dic455LP, em480/40). Fluorescence backgrounds were subtracted using the modal-values from the fluorescence images. Quantifications of cellular fluorescence were done using the ObjectJ plug-in of ImageJ^35^. Living cells were imaged directly from the growing cultures and fixed cells directly after fixation and PBS washing. After fixation the samples were allowed to mature overnight at RT, washed once more with 1 ml PBS and were then imaged. These samples are indicated in the text as “matured”.

### Western blot analysis

Samples for SDS–polyacrylamide gel electrophoresis and Western blotting were prepared by pelleting live cells and equalizing biomass in MilliQ and subsequent boiling (95 °C) in sample buffer buffer (0.0625 M Tris-Hcl pH 6.8, SDS 2 %, glycerol 10%, bromphenol blue 0.001 %) with and without dithiothreitol (0.1 M) reducing agent. Samples were loaded on 10 % SDS-PAGE gels (stacking buffer (0.5M Tris-HCl pH 6.8, 0.4 % SDS), separating buffer (1.5 M Tris-HCl pH 8.7, 0.4 % SDS), separated by electrophoresis and transferred on nitrocellulose membrane (Biorad) by wet blotting, bound antibodies were detected with the Odyssey FC scanner (LI-COR). The used antibodies were polyclonal rabbit α-GFP (1:2,000, Fisher Scientific) and polyclonal goat α-rabbit IRDye^®^ 680LT (1:20,000, LI-COR).

### FRET experiments and spectral measurements

FRET experiments were performed as described in^36^ with modifications for the sfTq2-mNG FRET pair. Acceptor and donor emission spectra were collected with a fluorometer (Photon Technology International, NJ) through 6 nm slit widths with 1 s integration time per scanned nm and 3 times averaging. For the acceptor (mNG) channel samples were excited by the monochromator set at 504 nm through a 500 ± 10 nm single band pass (BP) filter (BrightLine, Semrock, Rochester, NY) and emission wavelengths from 512 to 650 nm at 1 nm increments were measured through a 510 nm long-pass (LP) filter (Chroma technology corp., Bellow falls, VT). This spectrum was used to determine the amount of mNG in the sample. For the donor (mTq2, sfTq2 or sfTq2^ox^) channel samples were excited by the monochromator set at 450 nm through a 435 ± 40 nm BP filter (Semrock) and emission wavelengths from 470 to 650 nm at 1 nm increments were collected through a 458 nm LP filter (Semrock). Knowing the amount of mNG present in the sample and the shape of the mNG and mTq2 variant reference spectra and background fluorescence spectrum in the cells, the sample spectra were unmixed into their separate components: background fluorescence, mTq2, mNG and sensitized emission (FRET). The FRET efficiencies were calculated using the published algorithms^36,37^, using the spectral properties of mTq2 and mNG^8,9^. Spectral measurements using a multimode plate reader (BIOTEK Synergy MX, BioTek Instruments Inc., Winooski, VT) were performed as described ^4,36^ with the mTq2 variant donor channel acquisition set at 450 nm and emission scanning from 470 to 650 nm with minimal slit widths of 9 nm.

### Frequency domain fluorescence lifetime measurements

FLIM experiments were essentially performed as described before^8^. For periplasmic samples in solution, the exposure time was 500 ms, and the number of phase steps 18.

### OSER assay

The OSER assay uses an ER localization sequence (CytERM) that was inserted in the multiple cloning site of pmTurquoise2-N1 to obtain CytERM-mTurquoise2 (Addgene plasmid Plasmid #98833). The coding sequence of mTurquoise2 was replaced by the superfolder variants. After transfection of the plasmids, the localization of the fusion was visualized by confocal microscopy. The OSER assay was performed as described before^16^.

### Molecular brightness assay

The cellular brightness assay was performed in HeLa cells as described before^10^.

## References

1. Weiner, J. H. & Li, L. Proteome of the Escherichia coli envelope and technological challenges in membrane proteome analysis. Biochim. Biophys. Acta 1778, 1698–713 (2008).

2. Harvey, B. R. et al. Anchored periplasmic expression, a versatile technology for the isolation of high-affinity antibodies from Escherichia coli-expressed libraries. Proc. Natl. Acad. Sci. U. S. A. 101, 9193–8 (2004).

3. Piston, D. W. D. W. & Kremers, G.-J. G. J. Fluorescent protein FRET: the good, the bad and the ugly. Trends Biochem. Sci. (2007). doi:10.1016/j.tibs.2007.08.003

4. Meiresonne, N. Y., van der Ploeg, R., Hink, M. A. & den Blaauwen, T. Activity-related conformational changes in D,D-carboxypeptidases revealed by in vivo periplasmic förster resonance energy transfer assay in escherichia coli. MBio 8, e01089–17 (2017).

5. Costantini, L. M. et al. A palette of fluorescent proteins optimized for diverse cellular environments. Nat. Commun. 6, 7670 (2015).

6. Aronson, D. E., Costantini, L. M. & Snapp, E. L. Superfolder GFP is fluorescent in oxidizing environments when targeted via the Sec translocon. Traffic 12, 543–8 (2011).

7. Pédelacq, J.-D., Cabantous, S., Tran, T., Terwilliger, T. C. & Waldo, G. S. Engineering and characterization of a superfolder green fluorescent protein. Nat. Biotechnol. 24, 79–88 (2006).

8. Goedhart, J. et al. Structure-guided evolution of cyan fluorescent proteins towards a quantum yield of 93%. Nat. Commun. 3, 751 (2012).

9. Shaner, N. C. et al. A bright monomeric green fluorescent protein derived from Branchiostoma lanceolatum. Nat. Methods 10, 407–9 (2013).

10. Mastop, M. et al. Characterization of a spectrally diverse set of fluorescent proteins as FRET acceptors for mTurquoise2. Sci. Rep. 7, 11999 (2017).

11. Fukuda, H., Arai, M. & Kuwajima, K. Folding of green fluorescent protein and the cycle3 mutant. Biochemistry 39, 12025–32 (2000).

12. Cranfill, P.J. et al. Quantitative assessment of fluorescent proteins. Nat. Methods 13, 557–562 (2016).

13. Jain, R. K., Joyce, P. B., Molinete, M., Halban, P. A. & Gorr, S. U. Oligomerization of green fluorescent protein in the secretory pathway of endocrine cells. Biochem. J. 360, 645–9 (2001).

14. Goedhart, J. et al. Bright cyan fluorescent protein variants identified by fluorescence lifetime screening. Nat. Methods (2010). doi:10.1038/nmeth.1415

15. Kim, J. H. et al. High Cleavage Efficiency of a 2A Peptide Derived from Porcine Teschovirus-1 in Human Cell Lines, Zebrafish and Mice. PLoS One 6, e18556 (2011).

16. Costantini, L. M., Fossati, M., Francolini, M. & Snapp, E. L. Assessing the Tendency of Fluorescent Proteins to Oligomerize Under Physiologic Conditions. Traffic 13, 643–649 (2012).

17. den Blaauwen, T., Andreu, J. M. & Monasterio, O. Bacterial cell division proteins as antibiotic targets. Bioorg. Chem. 55, 27–38 (2014).

18. Liu, B., Persons, L., Lee, L. & de Boer, P. A. J. Roles for both FtsA and the FtsBLQ subcomplex in FtsN-stimulated cell constriction in Escherichia coli. Mol. Microbiol. 95, 945–970 (2015).

19. Robichon, C., Karimova, G., Beckwith, J. & Ladant, D. Role of leucine zipper motifs in association of the Escherichia coli cell division proteins FtsL and FtsB. J. Bacteriol. 193, 4988–4992 (2011).

20. Koushik, S. V., Blank, P. S. & Vogel, S. S. Anomalous Surplus Energy Transfer Observed with Multiple FRET Acceptors. PLoS One 4, e8031 (2009).

21. Steiert, F., Petrov, E. P., Schultz, P., Schwille, P. & Weidemann, T. Photophysical Behavior of mNeonGreen, an Evolutionarily Distant Green Fluorescent Protein. Biophys. J. 114, 2419–2431 (2018).

22. Karimova, G., Dautin, N. & Ladant, D. Interaction network among Escherichia coli membrane proteins involved in cell division as revealed by bacterial two-hybrid analysis. J. Bacteriol. 187, 2233–43 (2005).

23. Glas, M. et al. The soluble periplasmic domains of Escherichia coli cell division proteins FtsQ/FtsB/FtsL form a trimeric complex with submicromolar affinity. J. Biol. Chem. (2015). doi:10.1074/jbc.M115.654756

24. Condon, S. G. F. et al. The FtsLB subcomplex of the bacterial divisome is a tetramer with an uninterrupted FtsL helix linking the transmembrane and periplasmic regions. J. Biol. Chem. (2018). doi:10.1074/jbc.RA117.000426

25. Villanelo, F., Ordenes, A., Brunet, J., Lagos, R. & Monasterio, O. A model for the Escherichia coli FtsB/FtsL/FtsQ cell division complex. BMC Struct. Biol. (2011). doi:10.1186/1472-6807-11-28

26. Bart van den Berg van Saparoea, H. et al. Fine-mapping the contact sites of the Escherichia coli cell division proteins FtsB and FtsL on the FtsQ protein. J. Biol. Chem. 288, 24340–24350 27.(2013).

27. Kureisaite-Ciziene, D. et al. Structural analysis of the interaction between the bacterial cell division proteins FtsQ and FtsB. bioRxiv 362335 (2018). doi:10.1101/362335

28. van der Ploeg, R., Goudelis, S. T. & den Blaauwen, T. Validation of FRET Assay for the Screening of Growth Inhibitors of Escherichia coli Reveals Elongasome Assembly Dynamics. Int. J. Mol. Sci. 16, 17637–54 (2015).

29. Bethesda Research Laboratories. E. coli DH5 alpha competent cells. Focus - Bethesda Res. Lab. 8, 9 (1986).

30. Taschner, P. E., Huls, P. G., Pas, E. & Woldringh, C. L. Division behavior and shape changes in isogenic ftsZ, ftsQ, ftsA, pbpB, and ftsE cell division mutants of Escherichia coli during temperature shift experiments. J. Bacteriol. 170, 1533–40 (1988).

31. Denome, S. A., Elf, P. K., Henderson, T. A., Nelson, D. E. & Young, K. D. Escherichia coli mutants lacking all possible combinations of eight penicillin binding proteins: Viability, characteristics, and implications for peptidoglycan synthesis. J. Bacteriol. 181, 3981–3993 (1999).

32. Nørholm, M. H. A mutant Pfu DNA polymerase designed for advanced uracil-excision DNA engineering. BMC Biotechnol. 10, 21 (2010).

33. Koppelman, C.-M. et al. R174 of Escherichia coli FtsZ is involved in membrane interaction and protofilament bundling, and is essential for cell division. Mol. Microbiol. 51, 645–57 (2004).

34. Edelstein, A., Amodaj, N., Hoover, K., Vale, R. & Stuurman, N. Computer control of microscopes using manager. Current Protocols in Molecular Biology (2010). doi:10.1002/0471142727.mb1420s92

35. Vischer, N. O. E. et al. Cell age dependent concentration of Escherichia coli divisome proteins analyzed with ImageJ and ObjectJ. Front. Microbiol. 6, 586 (2015).

36. Meiresonne, N., Alexeeva, S., van der Ploeg, R. & den Blaauwen, T. Detection of Protein Interactions in the Cytoplasm and Periplasm of Escherichia coli by Förster Resonance Energy Transfer. Bio-Protocol 7, (2018).

37. Alexeeva, S., Gadella, T. W. J., Verheul, J., Verhoeven, G. S. & den Blaauwen, T. Direct interactions of early and late assembling division proteins in Escherichia coli cells resolved by FRET. Mol. Microbiol. 77, 384–98 (2010).

